# Identification of potential astrocytes in the teleost brain

**DOI:** 10.1101/2021.12.27.474298

**Authors:** Germaine Jia Hui Tan, Kathleen Wen Bei Cheow, May Si Min Ho, Suresh Jesuthasan

## Abstract

Astrocytes are abundant star-shaped glial cells in the mammalian brain, with essential roles in metabolism, development, homeostasis, response to injury, behavior, and learning. Surprisingly, most regions of the teleost brain are thought to lack astrocytes, based primarily on the use of GFAP (glial fibrillary acidic protein) as a marker^1^. Here, drawing on recent evidence that astrocytes are molecularly heterogeneous, we propose that astrocytes exist in the teleost brain, albeit of the *olig2* subtype^2^. Highly branched cells are present throughout the zebrafish brain, as shown here in *Tg(sox10:EGFP)* fish and previously in *Tg(olig2:GFP)* fish. Transcriptome data indicates the presence of brain cells that are *olig2* and *sox10* positive, which also express the astrocyte markers *sox9b, sparcl1* and *slc1a2b* but lack *gfap* and the oligodendrocyte marker *mbp*. In situ hybridization confirms that stellate sox10:EGFP cells express *olig2* and *sox9b*, while immunofluorescence indicates that they lack HuC/D and GFAP. We suggest that these cells be classified as astrocytes as this may more accurately reflect their functions.

## Main Text

Astrocytes were historically defined by their stellate morphology and more recently by molecular markers. One such marker is the glial fibrillary acidic protein (GFAP). Numerous attempts have failed to detect astrocytes in the forebrain and midbrain of teleosts^1^. Instead, radial glia with bushy processes were seen. Based on these findings, it has been proposed that zebrafish and other teleosts lack stellate astrocytes, and that astrocyte functions are performed by a subset of radial glia^3^. More recently, transgenesis with the promoter for *slc1a3b/GLAST*, a gene encoding a glutamate transporter that is found in a subset of GFAP-positive astrocytes, led to the identification of cells with bushy protrusions in the spinal cord^4^. These cells express astrocyte markers such as glutamine synthetase, tile with one another and have calcium transients in response to norepinephrine. They appear similar to mammalian astrocytes, although their cell bodies are located near the ventricle.

Studies in the mouse have established that astrocytes are heterogenous and transcriptionally diverse. They do not all express GFAP, although genes such as *sox9* are expressed in many subtypes^5^. One GFAP-negative subset in the forebrain expresses *olig2*^2^. Olig2 astrocytes occupy different territories compared to GFAP astrocytes, and are defined as astrocytes based on morphology, presence of markers such as s100ß, glutamine synthetase and Sox9, and lack of the myelin binding protein (MBP) ^2^. Although *olig2* has been thought to function in formation of oligodendrocytes, the finding that the *olig2* mutant has deficits in astrocytes^6^ indicates that this transcription factor has roles beyond differentiation of oligodendrocytes. One possibility is that it is expressed in a precursor that gives rise to both oligodendrocytes and astrocytes^7^. Intriguingly, *olig2* has previously been shown to be expressed broadly in the zebrafish telencephalon^8^. Only a minority (∼16%) of cells expressed MBP, and none expressed the neuronal marker HuC. Instead, virtually all (∼98%) co-expressed *sox10*. Strikingly, the *olig2*/*sox10* cells have multiple protrusions. Could some of these be astrocytes?

An examination of the single cell transcriptome of the zebrafish brain^9^ revealed the existence of two clusters (clusters 36 and 47) that are positive for *sox10* and *olig2* as well as other genes expressed in astrocytes, namely *s100ß* and *sox9*. One of these, Cluster 36, expressed *slc1a2b*, a glutamate transporter found in astrocytes, and *sparcl*, which codes for an ECM protein (Hevin) produced by astrocytes, but did not express *gfap* or *mbp*. Cluster 47, in contrast, expressed *mbp*. Cluster 36 is thus a candidate for an *olig2*-positive, *gfap*-negative astrocyte cluster.

Imaging of a *Tg(sox10:EGFP)* line revealed the existence of stellate cells that are broadly distributed throughout the brain (Fig 1A, B). In the optic tectum, cell bodies and branches were located throughout the neuropil. In the habenula, only one or two cells were detected per hemisphere, with branches extend throughout the structure. Similar cells were also observed in older fish, retaining their sparse distribution and extensive branching pattern. Stellate cells were not detected in 1 day-old fish; instead, radial cells were observed (Figure 1C). This suggests that *sox10* is expressed in progenitor cells at early stages of neural development, and then in stellate cells in the larval and mature brain. In situ hybridization confirmed that stellate EGFP-positive cells express *olig2* (Fig. 1D-F; 199/206 cells in 3 fish) and *sox9b* (Figure 1G-I; 69/79 cells in 3 fish), while immunofluorescence indicates that they lack HuC/D and GFAP (Fig. 1G-L; N = 3 fish each).

**Figure 1.**
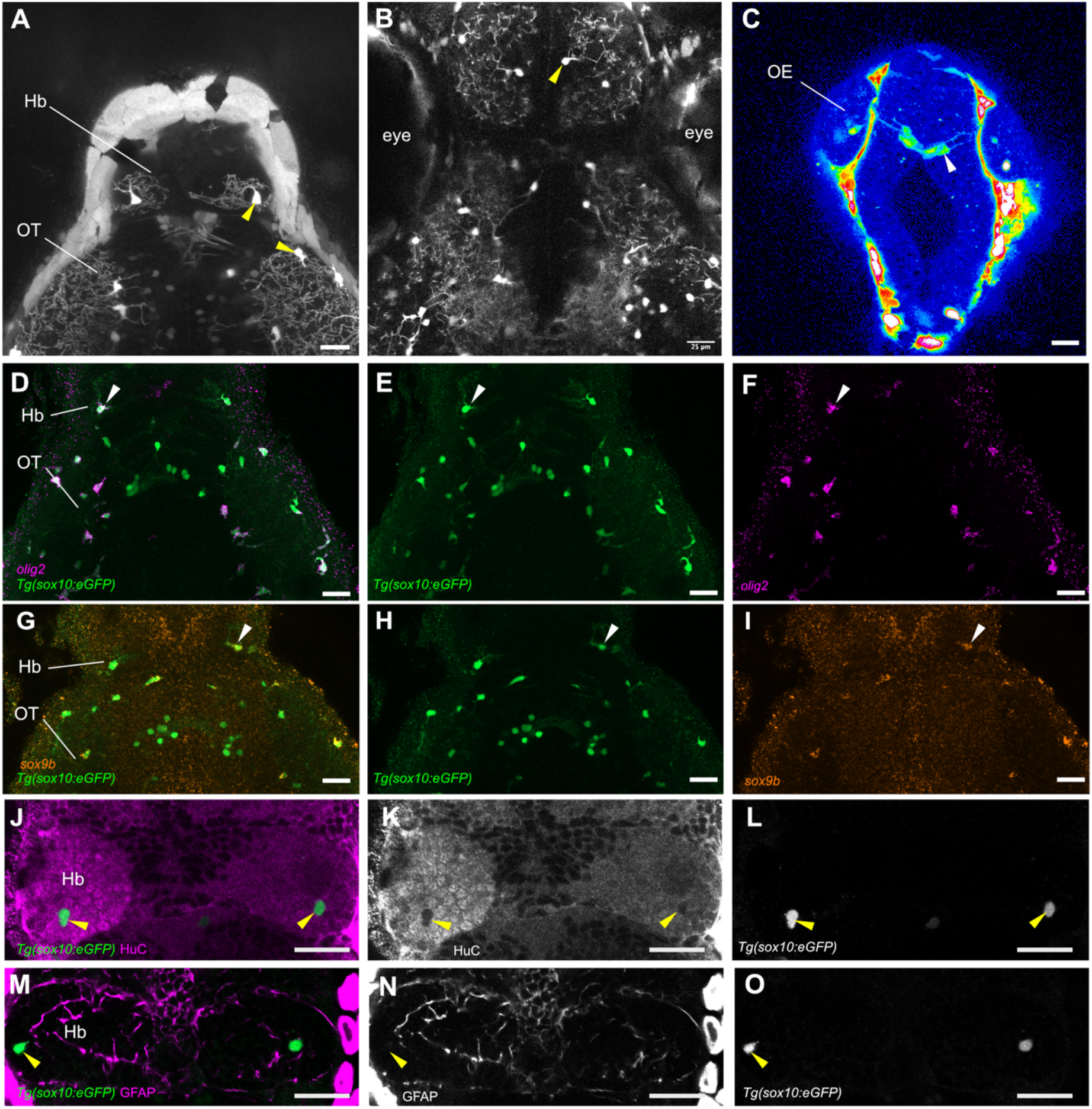
Characterization of branched cells in the brain of *Tg(sox10:EGFP)* fish. A-C. Imaging of larvae at 5 dpf (A) and 7 dpf(B), with GFP fluorescence in sparse cells (yellow arrowheads) and broadly distributed branches. At 1 dpf (C), cells with radial extensions were seen in the forebrain (white arrowhead). D-I. Expression of *olig2* (D-F) and *sox9b* (G-I) in 5 dpf Tg(sox10:EGFP) larvae, as detected by hybridization chain reaction. Arrowheads indicate EGFP-positive cells with co-expression. J-O. Antibody detection of the neuronal marker HuC/D (J-L) and GFAP (M-O). These images show the habenula of 6 dpf *Tg(sox10:EGFP)* fish. HuC/D and GFAP were not detected in GFP expressing cells (yellow arrowheads). All images are dorsal view, with anterior to the top. OE: olfactory epithelium; Hb: habenula; OT: optic tectum. Scale bar = 25 μm.

The cells described here appear identical to cells that have recently been termed oligodendrocyte progenitors^10^ or precursors. We suggest that astrocytes is a more accurate name, noting that the definition of what constitutes one cell type has a degree of subjectivity. Transcription factors, which are used to define cell type, have a role in determining what proteins are made, and thus what functions are possible. However, the genes that are transcribed depends on chromatin accessibility. Thus, in addition to transcription factors, morphology and function, which can be partially inferred from expression of genes mediating activities such as signaling, must also be considered. The *olig2*/*sox10* cells do not appear to be oligodendrocytes, as they largely lack the myelin binding protein^8^. While they may proliferate, we suggest that these cells do not function purely as progenitors, as they express genes associated with function such as *gad2, sparcl1* and *slc1a2b* (Table S1). A role as astrocytes, in contrast, is consistent with the expression of these genes. One may question the significance of naming these as astrocytes, rather than oligodendrocyte progenitors. We propose that nomenclature is important, as it influences thinking and shapes experiments. A cell type with features of astrocytes and *soxE* (a group that includes *sox9* and *sox10*) expression is present in basal vertebrates that lack myelinated axons^10^. An *olig2*/*sox10* cell in vertebrates may thus represent the progeny of an ancestral cell that fulfilled the critical functions of astrocytes before evolving the added ability to generate myelin in jawed vertebrates.

## Acknowledgements

This work was supported by the National Research Foundation (NRF2017-NRF-ISF002-2676). We thank Tom Carney for the *Tg(-4*.*9sox10:EGFP)*^*ba2*^ line and Anna Barron for discussions on glia.

## Supplemental Information

### Experimental Procedures

#### Fish

The *Tg(-4*.*9sox10:EGFP)*^ba2^ transgenic line, in which EGFP is driven by a sox10 promoter^1^, was maintained on a 14 hour:10 hour light/dark cycle in a Techniplast system. Embryos were kept in an incubator at 28°C, with a similar light/dark cycle. Experiments were performed under guidelines approved by the Institutional Animal Care and Use Committee of NTU (A19014 and A18013).

#### imaging of Tg(sox10:GFP) fish

Fish were anesthetized in buffered tricaine, then mounted in 2% low melting temperature agarose in E3 medium. Samples were imaged on a Zeiss LSM800 upright confocal microscope with a 40X water dipping objective.

#### In situ hybridisation

The *olig2* and *sox9b* qHCR probes were supplied by Molecular Instruments. Hybridization chain reaction^2^ was carried out according to the manufacturer’s HCR v3.0 protocol for whole mount zebrafish embryos and larvae. Labelled samples were imaged as above.

#### Antibody labelling

6 dpf *Tg(Sox10-eGFP)* fish were fixed overnight in 4% PFA at 4°C and washed with PBS before storage in 100% methanol at -20°C for 2 hours, followed by rehydration with reducing concentrations of methanol in 0.5% Triton x-100. Samples were treated with 10 to 20 ug/ml of proteinase K for an hour at room temperature on a tube rotator. This was followed by washing with PBS and blocking with 1% BSA and 0.1% triton X-100 in PBS for an hour at room temperature. Samples were incubated overnight at 4°C with the following primary antibodies – 1:50 anti-HuC/HuD mouse IgG2b (Thermo Fisher Scientific, MA, USA. Cat. No. A21271, Lot no. 2340692), 1:500 anti-GFAP polyclonal rabbit (Dako, Denmark. Cat. No. Z0334, Lot no. 41259205), 1:500 anti-GFP rabbit polyclonal (OriGene Technologies, MD, USA. Cat. No. TP401), and 1:500 anti-GFP mouse monoclonal (Thermo Fisher Scientific, MA, USA. Cat. No. A-11120). The secondary antibodies used were conjugated with Alexa Fluor 488 and 647 (Thermo Fisher Scientific), at a dilution of 1:1000. Samples were imaged as above.

## Supplemental Table

**Table S1:**
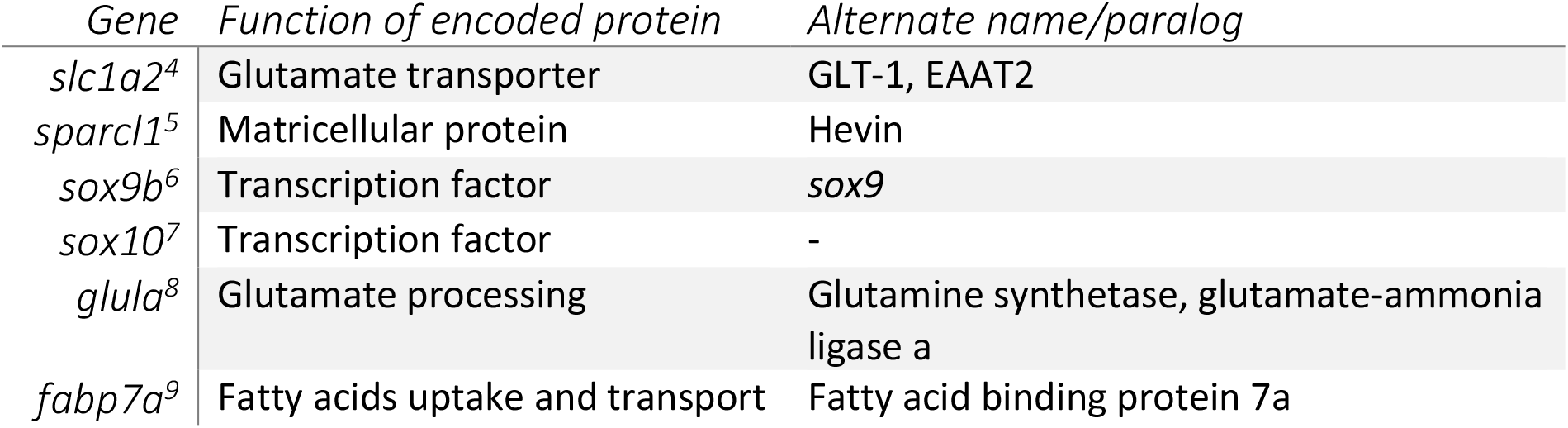
Genes in cluster 36^3^ that are associated with astrocytes in mammals.

## Supplemental Movie

**Movie S1**. (available at Figshare: 10.6084/m9.figshare.17696009). Z-stack of a 7 day old *Tg(-4*.*9sox10:EGFP)* fish. Planes are 0.72 μm apart. Gamma is set at 0.45.

## Notes

### Competing Interest Statement

The authors have declared no competing interest.

